# The use of DNA barcoding for the identification of giant hogweeds in the European North-East of Russia

**DOI:** 10.1101/2023.02.21.529251

**Authors:** D.M. Shadrin, I.V. Dalke, I.G. Zakhozhiy, D.S. Shilnikov, M.N. Kozhin, I.F. Chadin

## Abstract

Qualitative differences in the nrDNA ITS and ETS sequences in the total sample were determined to allow identification of the following groups of samples: *H. mantegazzianum, H*. cf. *sosnowskyi, H*. cf. *mantegazzianum* and *H*. cf. *sosnowskyi* × *H*. cf. *mantegazzianum* among other representatives of the *Heracleum* genus presented in the genetic databases. The study revealed the absence of correlation between a sample type in the group *H. mantegazzianum, H*. cf. *sosnowskyi, H*. cf. *mantegazzianum* and *H*. cf. *sosnowskyi* × *H*. cf. *mantegazzianum* and qualitative transformations in the ITS and ETS sequences. The qualitative analysis of the rbcL, matK, rps16 intron, intergenic spacers psbA-trnH, trnQ-rps16, rps16-trnK and rpl32-trnL cpDNA sequences for the samples identified as *Heracleum mantegazzianum*, *H*. cf. *sosnowskyi*, *H*. cf. *mantegazzianum* and *H*. cf. *sosnowskyi* × *H*. cf. *mantegazzianum* demonstrated no correlation between them as well.

## Introduction

Apiaceae Lindl. (Umbelliferae Juss.) is a large family of flowering plants comprised of compound species complexes with a high level of morphological variability (Kljuykov et al., 2018). The most popular classification for Umbelliferae at present is a system based on the anatomy and morphology of the fruit (Pimenov, Leonov, 1993; Pimenov, 2012; Valuyskikh et al., 2021). The classification system presented is not absolute and at times fails to accurately reflect relationships within the family (Downie et al., 2010). The genus *Heracleum* includes 53 taxa globally (Tkachenko, 2014). With regard to the *Heracleum* genus, most of the researchers’ attention is given to a group consisting of three species: *Heracleum mantegazzianum* Sommier & Levier, *H. sosnowskyi* Manden., *H. persicum* Desf. These three species are combined into a group of “large, tall or giant” hogweeds with a height of 3-5 meters (Mandenova, 1950, Nielsen et al., 2005; Jahodová et al., 2007a). Despite the fact that these species are referred to the section of the genus *Heracleum* – *Pubescentia* by a number of taxonomists, which includes some more species, allocating of the giant hogweed group is convenient for practical purposes. It is generally recognized that determination of the boundaries of the *Pubescentia* species is challenging (Mandenova, 1950; Jahodová et al., 2007a; Pimenov, Ostroumova, 2012; Ebel et al., 2018; Vladimirov et al., 2019).

Ornamental attributes and nutritional properties of these giant hogweed species caused their intentional introduction in a range of European countries. Following the second half of the 20th century, these species began to actively spread beyond the boundaries of cultivated crops. *H. mantegazzianum* Somm. et Lev. – Central and Western Europe (https://www.gbif.org/species/3034825), *H. persicum* Desf. ex Fisch., – European North (https://www.gbif.org/species/3628745) and *H. sosnowskyi* Manden – Eastern Europe (https://www.gbif.org/species/3642949. Currently, all the three species are considered invasive in a large area of the European continent, with the exception of the Caucasus, where *H. mantegazzianum* and *H. sosnowskyi* were originally described as endemic species (Nielsen et al., 2005; Ecology and management…, 2007; Zakhozhiy et al., 2021).

The *Heracleum* genus claims a complex evolutionary history, including hybridization, which resulted in complex variations in its species morphology, thus making identification of the species with traditional taxonomic methods more difficult (Pyšek et al., 2007; Sortland, 2010). Studies of the biology of the *H. mantegazzianum* and *H. sosnowskyi* species carried out within their invasive habitats in order to develop measures to control distribution of these species showed that they can be considered ecological twins for a number of individual development, structural and population dynamic features (Pyšek et al., 2007; Dalke et al., 2015). It should also be taken into account that their identification and documentation were not always carried out when introducing plants of this genus (Kudinov et al., 1980; Skupchenko, 1989; Pyšek et al., 2007). According to some authors (Pyšek et al., 2007), it would be extremely difficult for introducers to collect mericarps of solely one species in the common growth area for both *H. mantegazzianum* and *H. sosnowskyi*, since the key distinction feature of these two species – the degree of dissection of leaf lobes – is almost impossible to be used in the fruiting time. Some authors (Ebel et al., 2018) noted that “…the use of the name *Heracleum sosnowskyi* for wildings of the “giant hogweed” is rather tentative”. We believe that the statement about the conditionality of such naming can be equally applied to *H. mantegazzianum*.

In the last decade, in order to identify systematically complex plants, alongside with traditional taxonomy methods DNA barcoding of plants has been widely used, which enables identification of organisms based on short fragments of the DNA sequence (Hebert et al., 2003; Kress, 2017). In order to barcode representatives of the plant kingdom the following is used: *rbc*L and *mat*K genes, intergenic spacer *trn*H-*psb*A of chloroplast DNA (cpDNA), sequence ITS1-5.8s-ITS2 of the ribosomal nuclear DNA cluster (nrDNA) (Shneyer, Rodionov, 2018; Shekhovtsov et al., 2019; Shadrin, 2021). By now, the most detailed description of the molecular phylogenetic relationships, as well as the origin and migration of representatives of the Apiaceae family has been carried out using the ITS region of nrDNA (Downie et al., 2001, 2010; Banasiak et al., 2013). To identify a number of subtaxa of the Apiaceae family, an attempt was made to use the cpDNA *trn*H-*psb*A intergenic spacer region (Degtjareva et al., 2012), as well as a combination of ITS and *trn*H-*psb*A sequences (Liu et al., 2014; Valuyskikh, Shadrin, 2021). Another work declared high conservation of the intergenic spacer *psb*A*-trn*H to act as a limitation for the identification of representatives of the *Heracleum* genus and related taxa (Logacheva et al., 2007). An effective marker for phylogenetic studies of the tribe Tordylieae, which comprises the *Heracleum* genus, is the nrDNA ETS (external transcribed spacer) sequence, especially in combination with the ITS sequence (Logacheva et al., 2010). In addition to the markers listed above, for the molecular phylogenetic analysis of representatives of the *Heracleum* genus in the Chinese flora, noncoding cpDNA sequences were used: rps16 intron, trnQ-rps16, rps16-trnK and rpl32- trnL (Yu et al., 2011). The discrepancy between the phylogenies of the *Heracleum* genus based on the comparison of nrDNA and cpDNA sequences was also noted there, which, according to the authors, is an unprecedented case in the molecular phylogenetic system of the Apiaceae system (Yu et al., 2011). The Apiaceae is a large and taxonomically complex family, especially at the low systematic level, which requires new approaches for the analysis and search for taxonomic attributes.

The work aims at finding cpDNA and nrDNA markers that can make it possible to differentiate between *H. sosnowskyi* and *H. mantegazzianum*, which is essential to solve the taxonomic problems for this group of plants and assess the vector and rate of microevolutionary processes of these species under the invasion conditions.

## Materials and methods

### Sampling

The samples of the Heracleum genus representatives were collected in five Russian regions (Fig. 1.1, 1.2): the Kabardino-Balkarian Republic (5 samples), Karachay-Cherkess Republic (14 samples), Komi Republic (16 samples), Krasnodar Krai (11 samples), Murmansk Region (4 samples). The samples were taken from reproductive plants in the flowering or initial fruiting phase. Samples dedicated for genetic analysis were put into paper bags. Herbarium specimens were obtained from each sample: a small segment of the inflorescence and the middle leaf cut into several parts according to the size of the herbarium sheet. Each plant was photographed from several angles, the height and diameter at the base were measured.

**Fig. 1.**
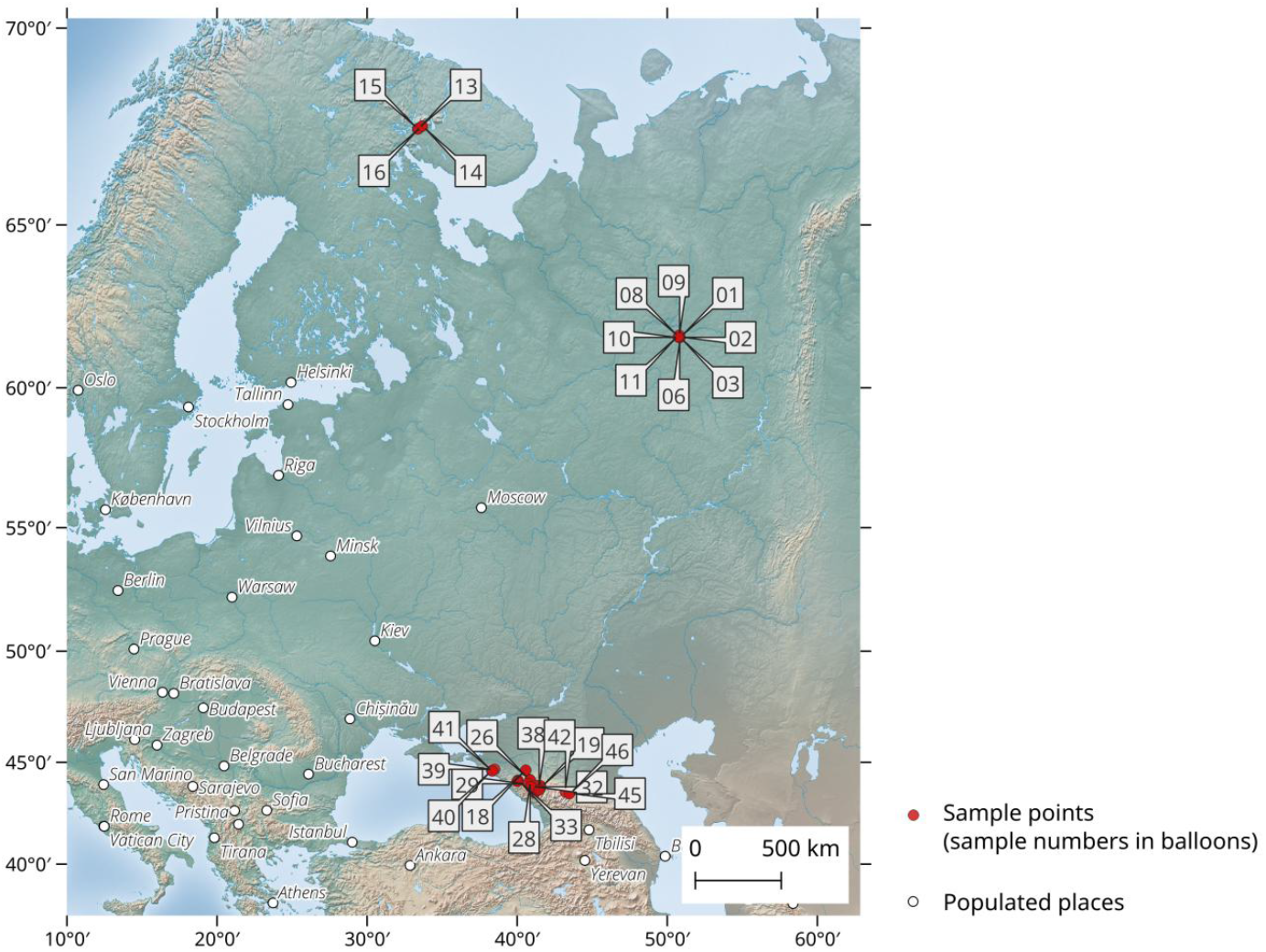
*Heracleum* samples collection sites within invasive and natural habitats.

**Fig. 2.**
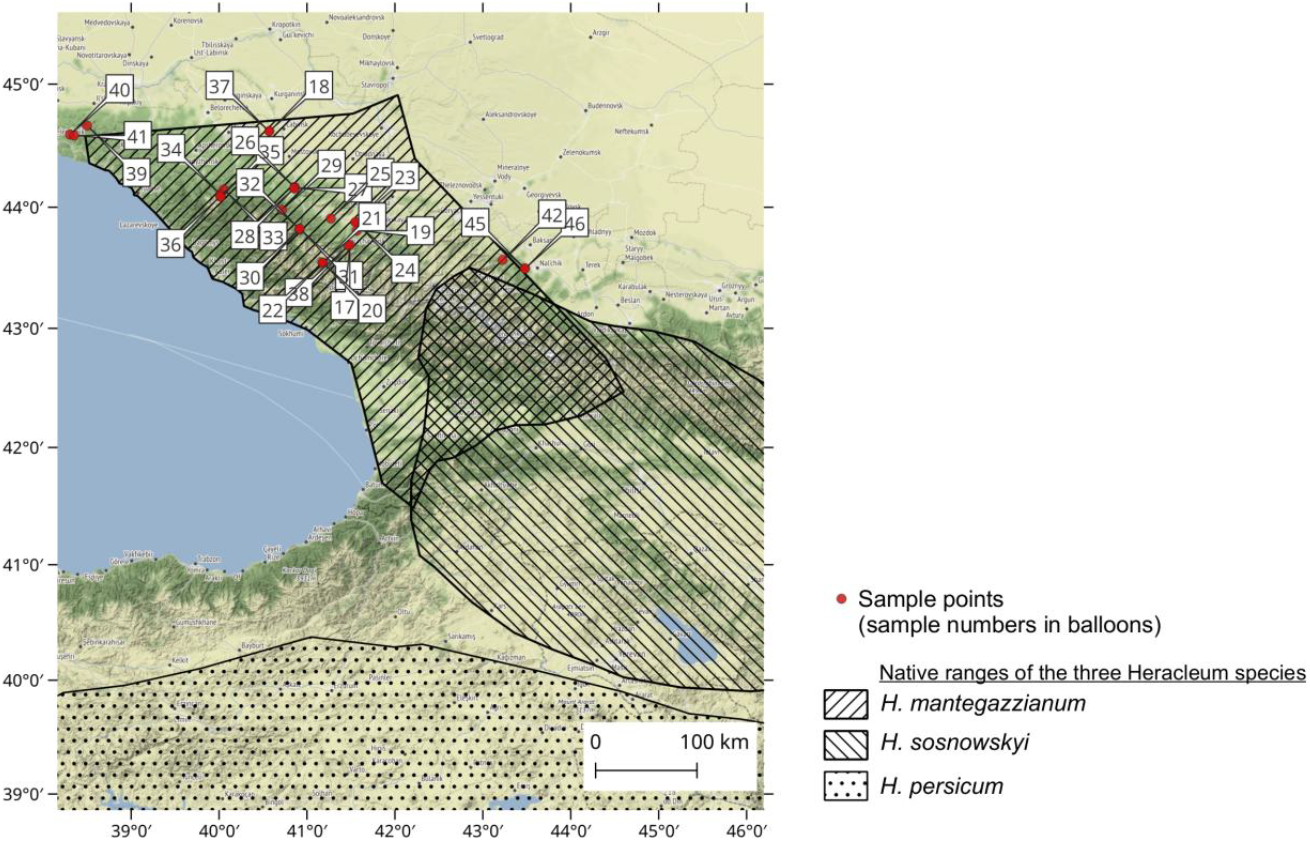
Boundaries of natural habitats for *Heracleum mantegazzianum*, *H. sosnowsky*, *H. persicum* and sample collection sites. Habitat boundaries are given according to Jahodová et al., 2007a.

### Morphological analysis

Samples of *Heracleum sibiricum* L. For the molecular phylogenetic analysis were identified using the key published in *«Flora of the Northeast of the European Part of the USSR»* (Flora…, 1977). We analyzed the attributes differentiating *H. mantegazzianum* and *H. sosnowskyi*, based on the data of a number of authors who had an opportunity to work with both species (Table 1) (Mandenova, 1950; Tsvelev, 2000; Pimenov, Ostroumova, 2012; Vladimirov et al., 2019). In most cases, the quantitative characteristics given for both species are overlapping sets. An exception is the plant height (up to 150 cm) indicated for *H. sosnowskyi*, according to I.P. Mandenova’s monograph (Mandenova, 1950). On this basis, all the giant hogweed samples we collected could not be referred to *H. sosnowskyi*. The most striking difference between these two species, as noted by all the authors, is the shape of segments and leaf lobes. I.P. Mandenova also points out the difference in leaf shape between *H. mantegazzianum* and *H. sosnowskyi*, however, she mentions that the leaf shape in plants of this genus is very variable and cannot serve as a stable diagnostic property. Other authors (Tsvelev, 2000; Vladimirov et al., 2019) consider features associated with leaf shape to be sufficient to distinguish between these two species.

**Table 1.** Attributes distinguishing *H. sosnowskyi* from *H. mantegazzianum*

| Author | Attribute | Significance for<br>*H. mantegazzianum* | Significance for<br>*H. sosnowskyi* |
| --- | --- | --- | --- |
| Mandenova I.P,<br>1950 | height (up to) | 300 cm | 150 cm |
|  | pubescence of lower part of leaf | scatteredly hairy | slightly protuberant |
|  | segment dissection | profoundly pinnatiparite | slightly pinnatiparite or trilobate |
|  | shape of segment lobes | triangular lanceolate, elongated towards the top, sharply pointed | wide egg-shaped |
|  | dissection of lobes of end segment | pinnate | slightly pinnate |
|  | shape of lobes of end segment | elongated and pointed | egg-shaped |
|  | colour of anther | pale olive | dark purple |
|  | shape of disc | torulose, conical | torulose, hemispheric |
| Pimenov M.G,<br>Ostroumova T.A.,<br>2012 | shape of leaf segments | egg-shaped or elongated egg-shaped, unequal-edged | wide egg-shaped |
|  | type of dissection of leaf segments | pinnatiparite | slightly 3-5-partite |
|  | shape of lobes of leaf segments | lanceolate, elongated towards the top, sharply pointed | sharp, but not long and pointed |
|  | type of dissection of lobes of end segment | profoundly pinnatiparite | tripartite |
|  | shape of lobes of end segment | lanceolate, pointed at the top | rounded |
|  | pubescence of umbel spokes | hairy, long (0.2-0.5 mm) hairs | puberulent |
|  | colour of anther | grey-green | grayish-yellow |
|  | shape of stylopodium (cone) | conical, torulose and wrinkled | laticonic |
|  | length of stylodium (stiles), mm | from 2 to 2.5 | from 1.5 to 2.0 |
| Vladimirov V. et al.,<br>2019 | shape of end lobes of lower and middle stem leaves | triangular or elongated triangular | short, egg-shaped or elliptical |
|  | distance between leaf lobes | almost equal to the width of leaf lobes | lobes touch or almost touch each other |
|  | shape of terminal lobe of terminal leaf segment | Rachises are thin-winged. At the base, elongated and wedge-shaped (length of the base is 2-5 times greater than the width) | Rachises with wide pteroid outgrowths. Not wedge-shaped at base (up to two times longer than the width) |
|  | presence of spines on small protuberances on the pteroid ribs of the mericarp | Absent | with sparse upturned small spikes |
| Tsvelev H.H., 2000 | degree of dissection of lobes of first order leaves | pinnatiparite | wider, pinnate or serrated |

For the morphological identification of the giant hogweed samples described in this work, we used the set of features given in the monograph by M.G. Pimenov and T.A. Ostroumova, 2012 (Pimenov, Ostroumova, 2012) (Table 1). These authors recognize two species – *H. sosnowskyi* and *H. mantegazzianum*, but for the latter one they consider *H. wilhemsii* Fissch. & Avé-Lall to be a preferable name, though they accept the use of *H. mantegazzianum* due to its being widely distributed. All the specimens collected in the Karachay-Cherkess Republic and Krasnodar Krai were associated by us with *H. mantegazzianum* based on morphological features, which is also consistent with the known boundaries of the natural habitats of the giant hogweed (Jahodová et al., 2007a) (Fig. 1). The identification was carried out mainly on the basis of the shape of the segments and leaf lobes of the samples. The use of the shape of the stylopodium, size of the stiles, and color of the anthers as attributes was challenged either due to the difficulties in correlating the verbal description of the morphology of these structures with the observed condition of the attributes, or because the observed condition of such attributes failed to correspond to any one described in the species diagnosis. In some cases, not only quantitative, but also qualitative properties turned out to be midpoint between the attributes typical for *H. sosnowskyi* and *H. mantegazzianum*. Each specimen of the giant hogweed was assigned to either *H. sosnowskyi* or *H. mantegazzianum* based on a count of the number of attributes that matched the description of the species. With an equal number of attributes in the sample assigned to each of these two species or with a predominance of intermediary characteristics, such samples were considered hybrids of *H. sosnowskyi* x *H. mantegazzianum*.

### Molecular Analysis

#### DNA Extraction, Amplification, Sequencing

Total DNA was extracted from dried leaves using the DNA-Extran-3 kit (Sintol, Russia) according to the manufacturer’s manual. Polymerase chain reaction (PCR) of the fragments was carried out in 50 μl of a mixture containing 10 μl of ScreenMix (Eurogen, Russia), 10 μl of each primer (0.3 μM) (Evrogen, Russia), 18 μl of ddH2O (PanEko, Russia) and 2 μl of DNA nanomatrix (1 ÷ 100 ng). Primers for the amplification of DNA sections used in the analysis and the conditions for carrying out the polymerase chain reaction are presented in Table 2.

**Table 2.**
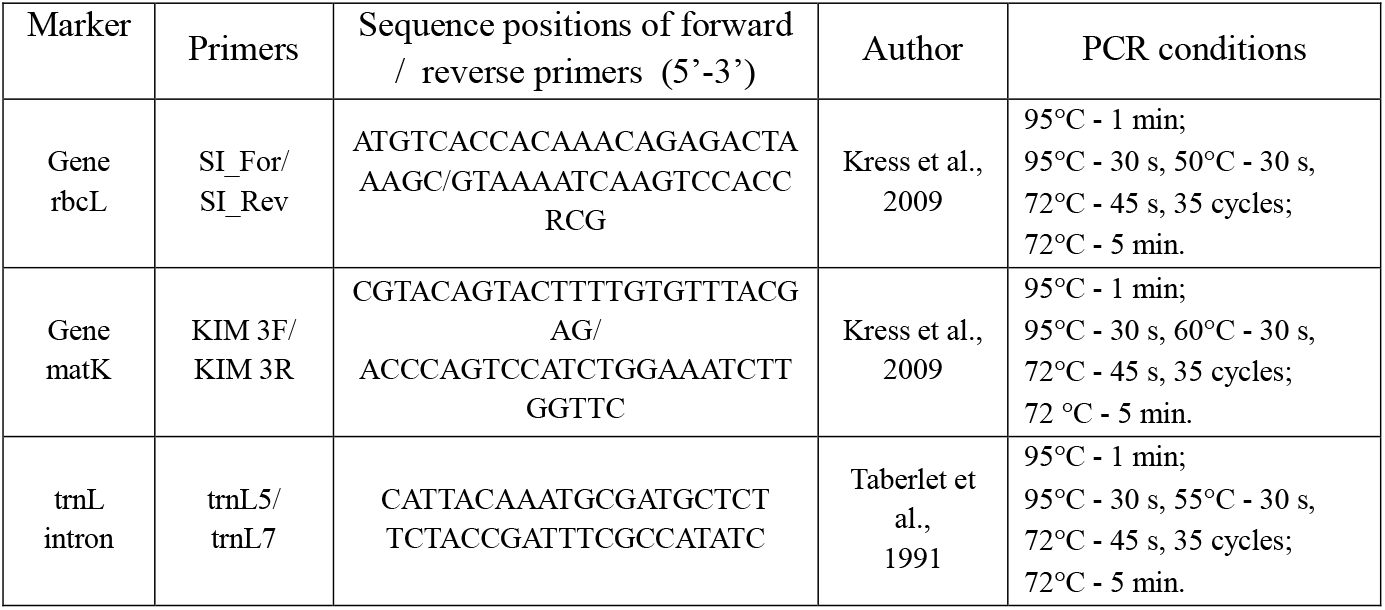
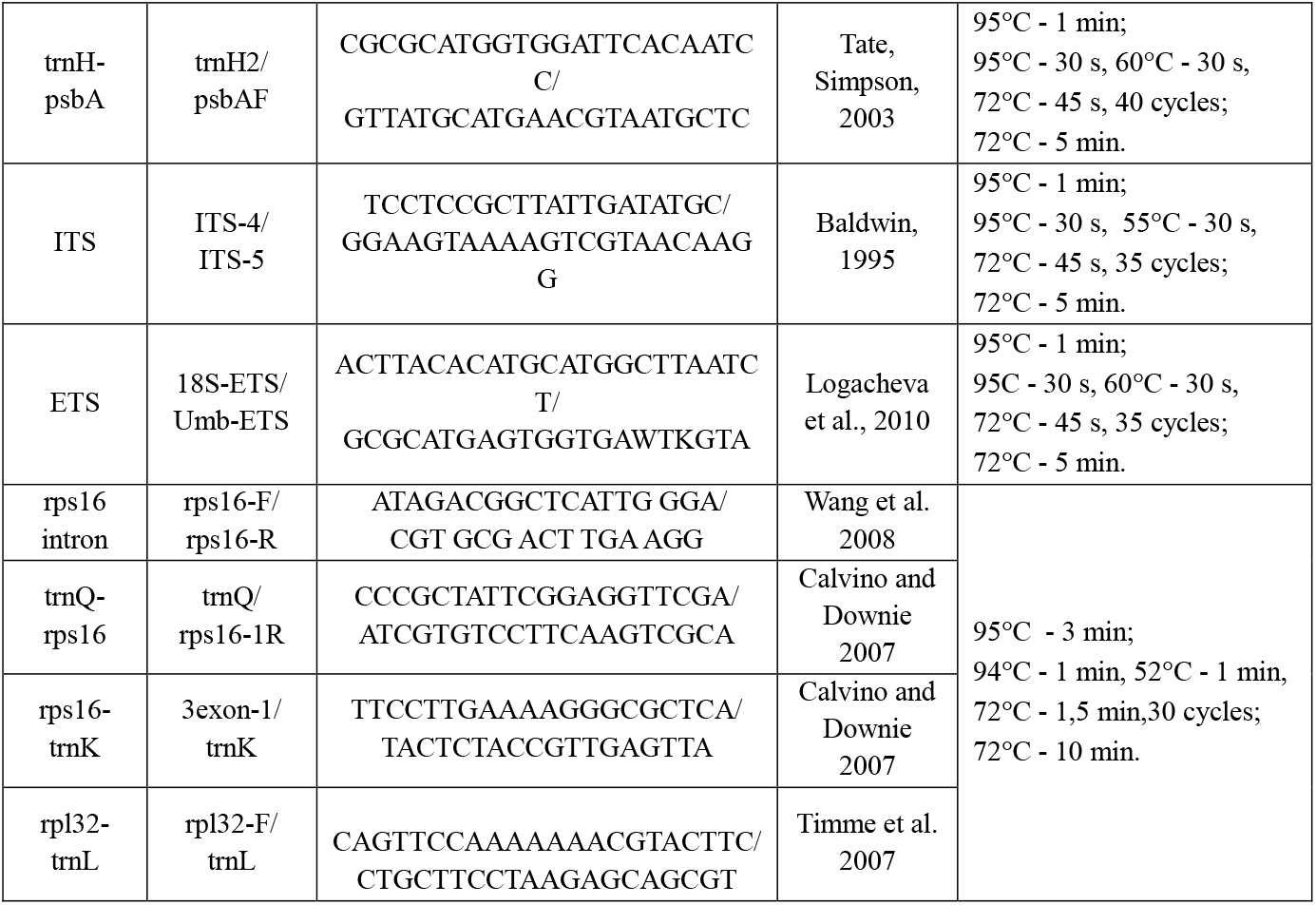
Used primers and conditions for the polymerase chain reaction

The PCR products were separated in a 1.5% agarose gel based on tris-acetate-ethylene-diamine-tetraacetic acid (TAE) and further purified using the ColGen DNA purification system (Sintol, Russia) according to the manufacturer’s manual. The sequencing products were separated on a NANOFOR 05 DNA sequencer (Sintol, Russia). DNA isolation, PCR, and sequencing were performed using the equipment of the “Molecular Biology” Center for Collective Use, the Institute of Biology, Federal Research Center of the Komi Research Center, Ural Branch of the Russian Academy of Sciences, Syktyvkar.

Multiple alignment of nucleotide sequences was performed using the ClustalW algorithm in MegaX (Thompson et al., 1994; Kumar et al., 2018). Phylogenetic trees were built in MegaX using the maximum likelihood method (Maximum Likelihood, ML). When constructing molecular phylogenetic trees, we used the nucleotide sequences obtained by us, as well as data from other authors available in Genbank (NCBI, 2023) and BOLD Systems (BOLD Systems, 2023) databases. *Azilia eryngioides* (Pau) Hedge & Lamond was used as an outgroup in the reconstruction of phylogenetic trees. The analysis resulted in obtainment of characteristics for both individual sequences and combined data. The list of samples of Heracleum genus considered in this study, including their location, voucher information, and GenBank (NCBI) number, is presented in Table 3.

**Table 3.**
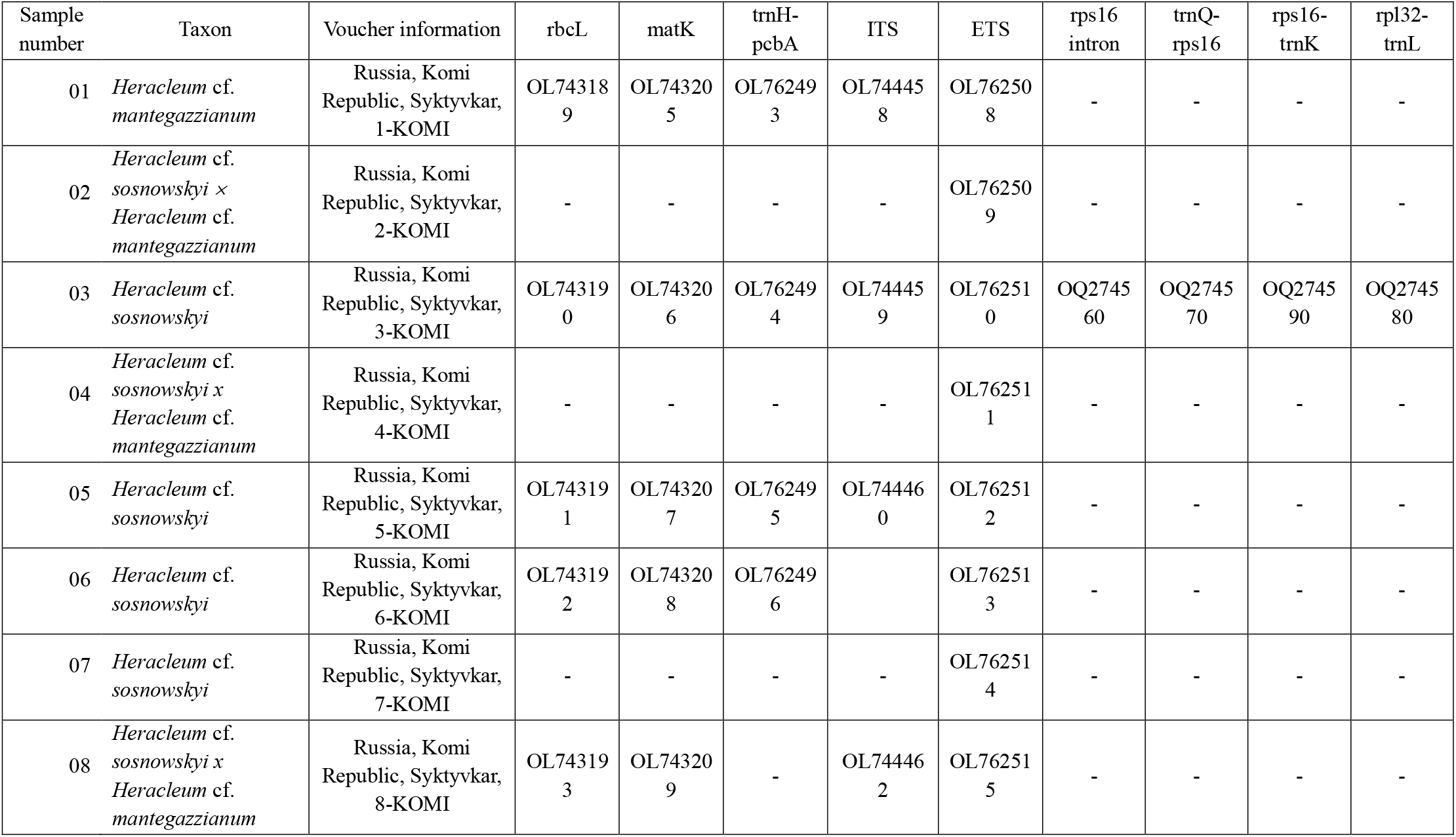
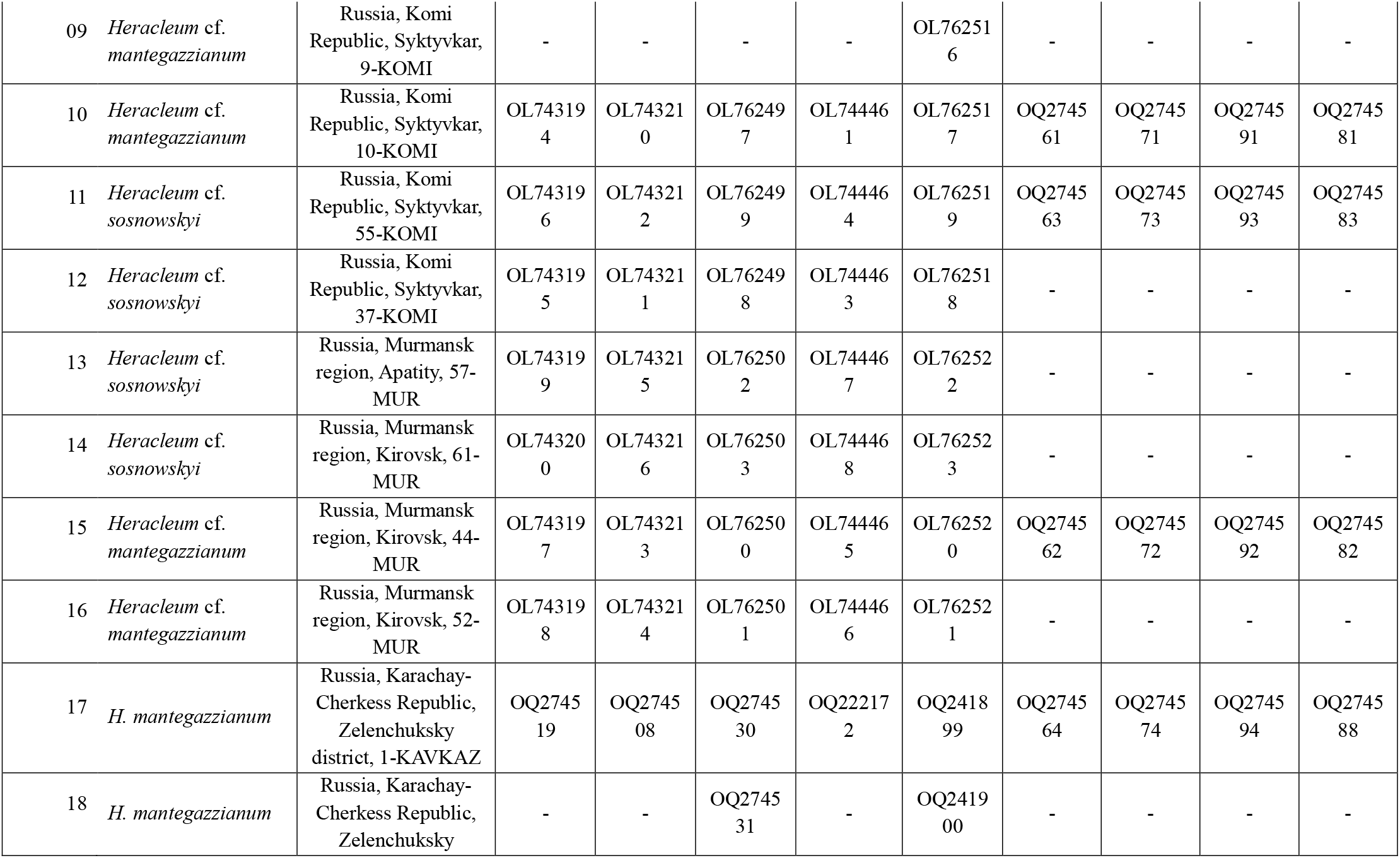
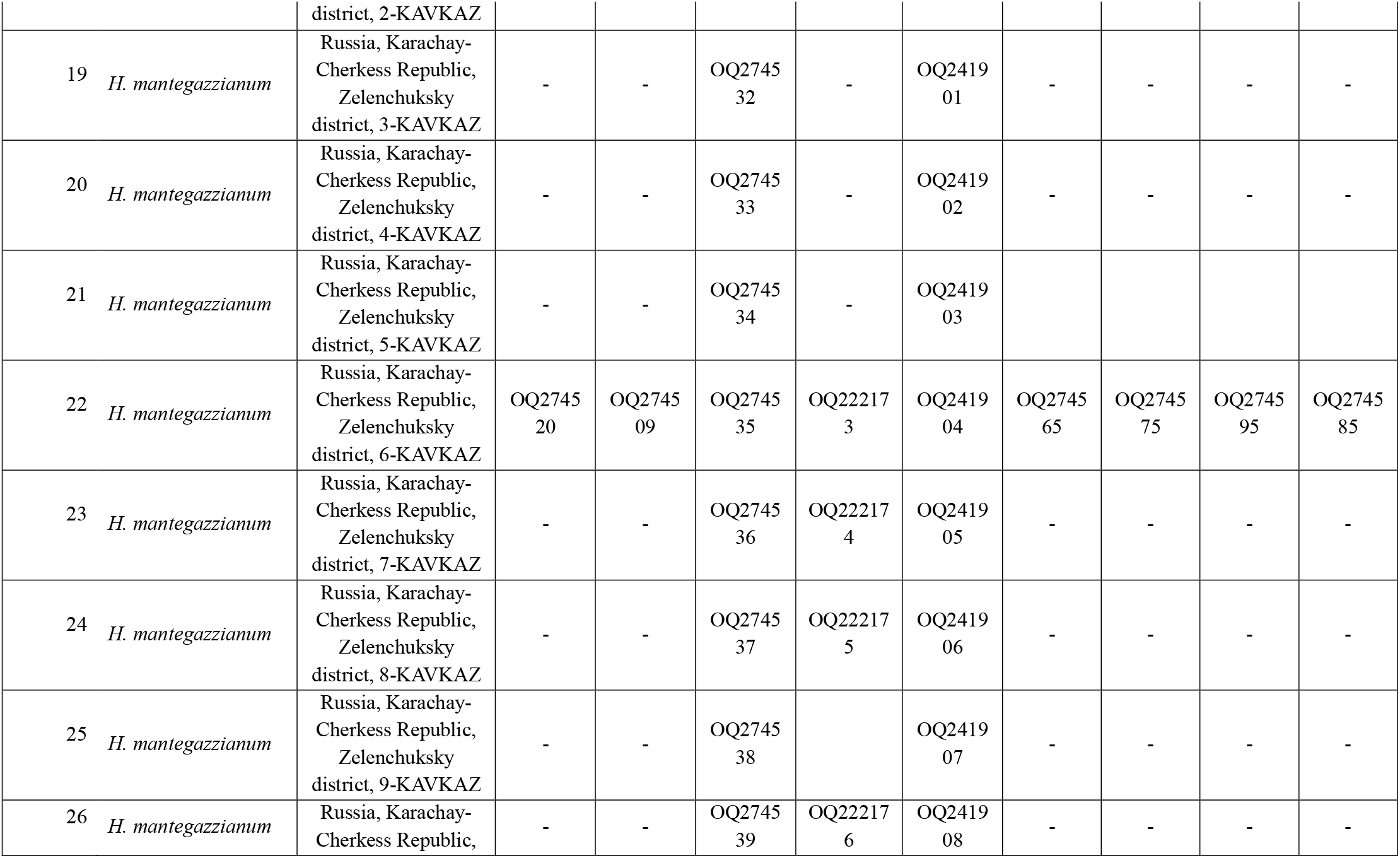
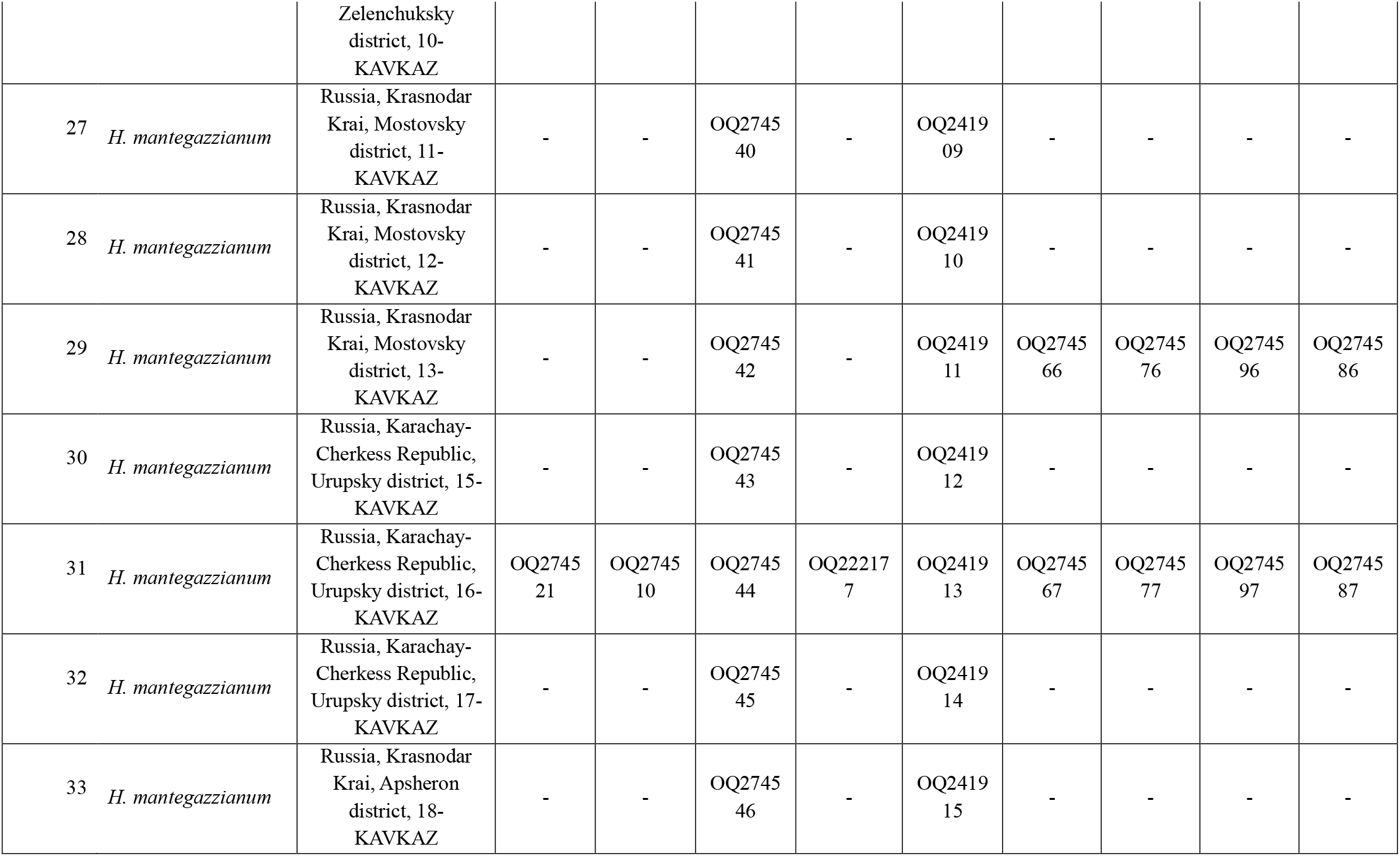
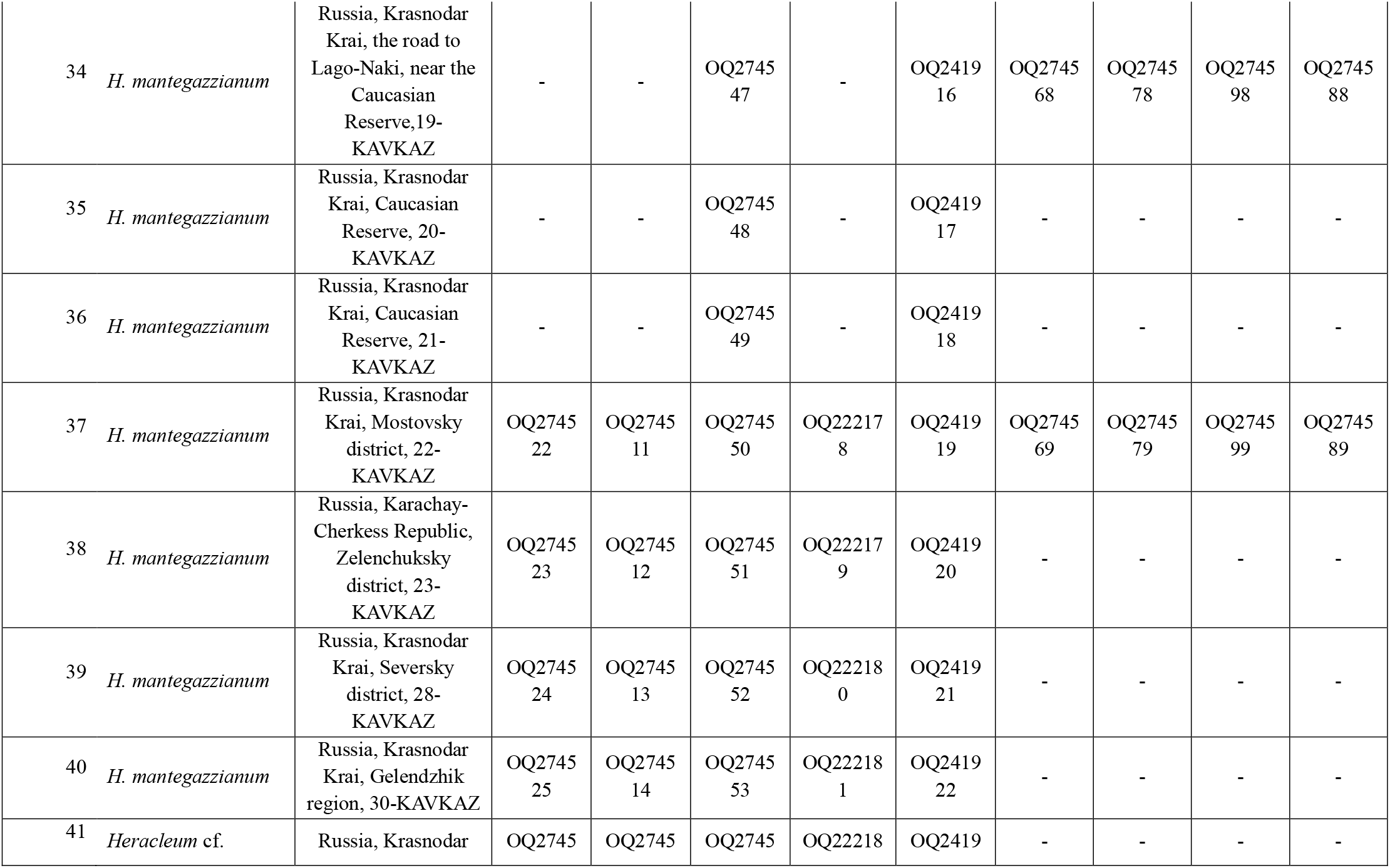
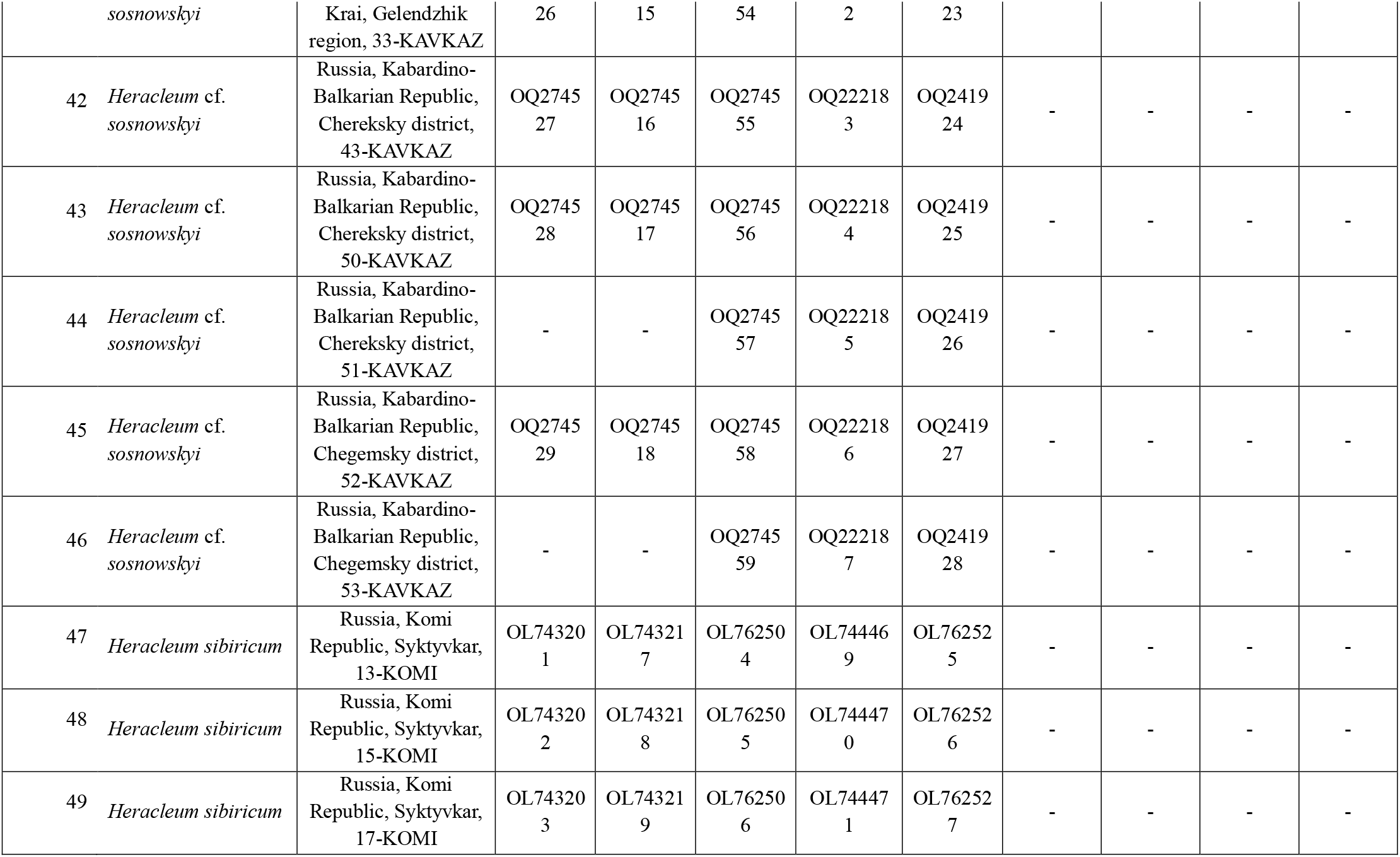
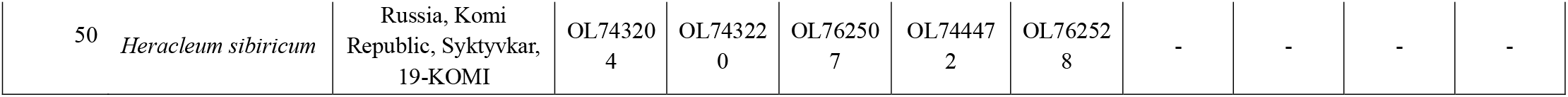
Origin, voucher information (or references), and GenBank accession numbers for plants used in the present study

## Results and Discussion

The molecular phylogenetic analysis included 50 samples of plants of the *Heracleum* genus (Table 3). DNA sequences of markers were not obtained for all the samples included in the analysis, since originally the analysis for each marker was carried out on a selective group that included plants classified by us as *H. sosnowskyi* and *H. mantegazzianum*. If the specimens of the sample group did not differ or produces differences not correlating with species specificity, the marker was not used further. A total of 224 sequences were generated for nine markers (Table 4).

**Table 4.** Characteristics of sequences of *Heracleum mantegazzianum*, *H*. cf. *sosnowskyi*, *H*. cf. *mantegazzianum and H*.cf. *sosnowskyi* × *H*.cf. *mantegazzianum*.

| Characteristic | rbcL | matK | psbA-trnH | rps16 intron | trnQ-rps16 | rps16-trnK | rpl32-trnL | ITS | ETS | Combined (ITS+ETS) |
| --- | --- | --- | --- | --- | --- | --- | --- | --- | --- | --- |
| Sequence length (bp) | 544 | 719 | 188-189 | 755-757 | 1217-1219 | 668-671 | 955-968 | 602 | 382-385 | 984-987 |
| Alignment positions | 544 | 719 | 189 | 757 | 1221 | 673 | 978 | 602 | 385 | 987 |
| Number of constant positions | 544 | 718 | 177 | 752 | 1210 | 667 | 970 | 600 | 369 | 969 |
| Number of variable characters | 0 | 1 | 11 | 3 | 8 | 1 | 7 | 2 | 16 | 18 |
| Number of parsimony-informative positions | 0 | 1 | 2 | 2 | 1 | 0 | 5 | 0 | 12 | 12 |
| Alignment positions/<br>Number of variable characters, % | 0 | 0,1 | 5,8 | 0,4 | 0,1 | 0,8 | 0,7 | 0,3 | 4,2 | 1,8 |

Based on the analysis of the published sequences of the *rbc*L, *mat*K genes and cpDNA *psb*A-*trn*H intergenic spacer of the plants belonging to the *Heracleum* genus (NCBI, 2023; BOLD Systems, 2023), it was assumed that the studied sequences are highly conserved and cannot be used for DNA barcoding of hogweed plants. Sequence analysis of the rbcL and matK genes of 27 samples collected by us (Table 3) jointly with other data for other species of the *Heracleum* genus (NCBI, 2023; BOLD Systems, 2023), showed a very low level of divergence both within the analyzed group of giant hogweeds and for the entire series of *Heracleum* taxa. The degree of pairwise genetic differences between the compared samples was from 0 to 1.1% and from 0 to 1.5% for the *rbc*L and *mat*K genes, respectively. Among the plant samples assigned by us to *H. sosnowskyi* and *H. mantegazzianum*, the degree of pairwise genetic differences was 0%. Thus, it was confirmed that these markers are not applicable for identification of plants of the *Heracleum* genus.

The analysis of 41 sequences of the *psb*A*-trn*H intergenic spacer revealed variations in patterns TTGTTCTATCA and TGATAGAACAA at positions from 60 to 76 bp, which were repeated twice in the analyzed sequences (Fig. 3). Previously, discrete variations in the *psb*A-*trn*H sequence were published for plants of the *Heracleum* genus (Logacheva et al., 2007).

**Fig. 3.**
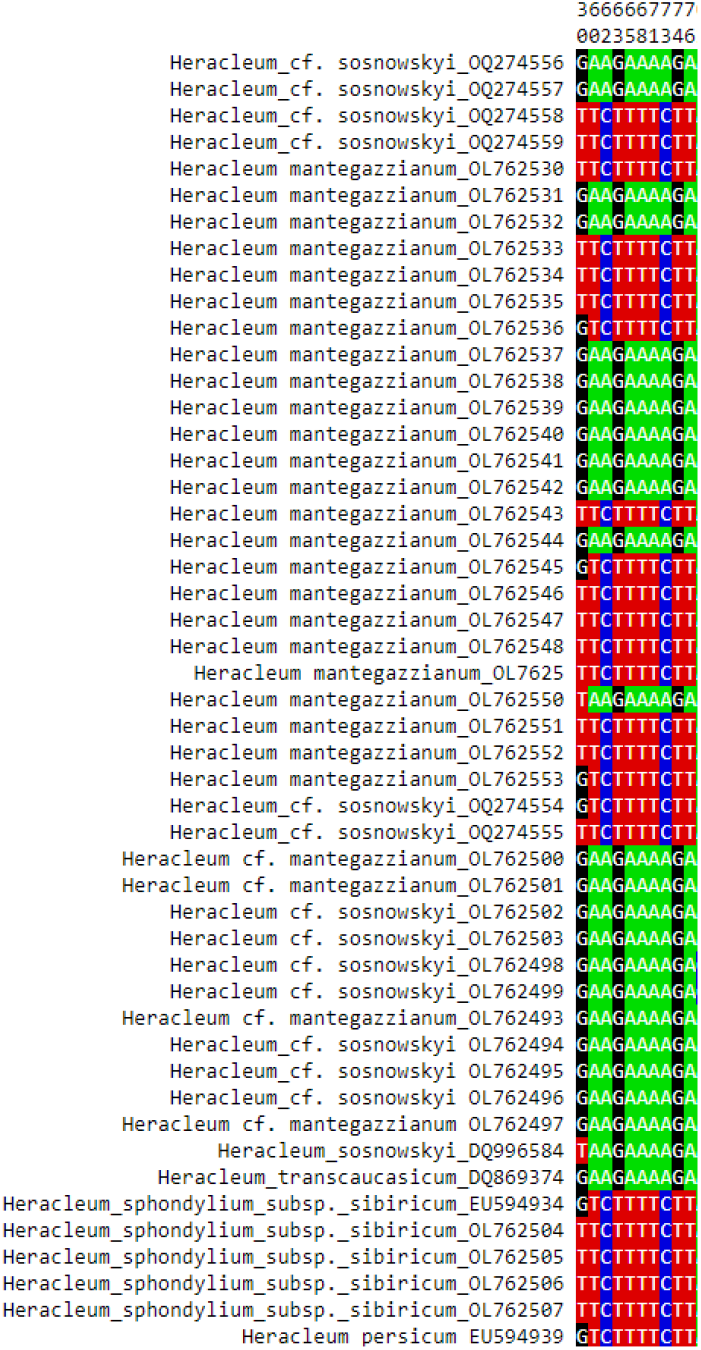
The variable sites from an alignment of *trn*H-*psb*A cpDNA sequences.

The variations described above occur both within our giant hogweed sample series and among other species included in the analysis. The analysis of variants showed no correlation between this type of variability and species. The presence of variations was explained by a combination of two independent evolutionary events (inversion and duplication), assuming that they occurred several times independently of each other (Logacheva et al., 2007). Following the *psb*A*-trn*H intergenic spacer sequence alignment, the studied sample series showed another polymorphic site – at position 30 (G/T). The revealed polymorphism, like the previous ones, did not correlate with the species. Pairwise genetic variability in the sample analyzed was 5.3%.

Next, 10 plant samples were studied to assess the variability of the sequences of rps16 intron and three non-coding cpDNA intergenic spacers – *trn*Q-*rps*16, *rps*16-*trn*K and *rpl*32-*trn*L (Table 3.1). Previously, these markers were used to study molecular phylogenetic relationships in the genus *Heracleum* (Yu et al., 2011) and to identify *H. mantegazzianum* (Identification…, 2019). Sequence alignment showed that rps16 intron and intergene spacer *trnQ-rps16* have a less pronounced level of variability than intergenic spacers *rps*16-*trn*K and *rpl*32-*trn*L (Table 4). The polymorphism of the rps16 intron fragments and the *trn*Q-*rps*16 intergenic spacer is mainly explained by single nucleotide substitutions. The variability of the *rps*16-*trn*K intergenic spacer was determined by single nucleotide insertions and deletions. In the hogweed samples under study, the *rpl*32-*trn*L intergenic spacer, which included three indelia 12, 10, and 6 bp long, with a total alignment length of 978 bp attracted special attention. For other angiosperms, specific indeliums were described in the sequence of the intergenic spacer *rpl*32-*trn*L (Shaw et al., 2007; Miller et al., 2009).

Nevertheless, the revealed variability for the rps16 intron sequences and cpDNA intergenic spacers trnQ-rps16, rps16-trnK and rpl32-trnL of the sample series under study did not allow to allocate the samples based on their reference to *H. mantegazzianum, H*. cf. *sosnowskyi* or *H*. cf. *mantegazzianum* (Fig. 4).

**Fig. 4.**
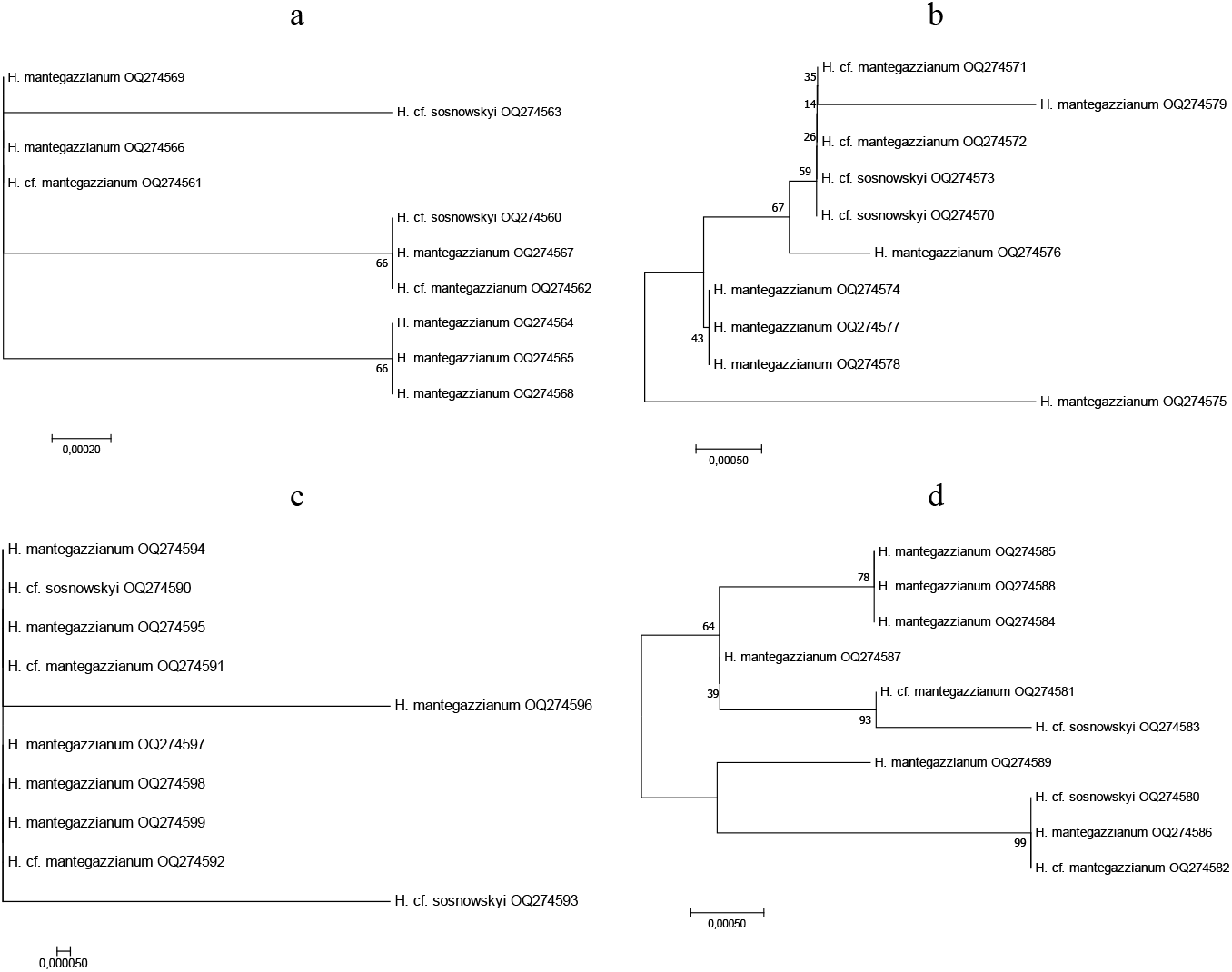
Molecular-phylogenetic trees constructed based on sequence comparison of 10 plant samples of giant hogweed. Trees were constructed using the maximum likelihood method. The specimen GenBank accession numbers in parentheses follows the names. a) - rps16 intron, b) - trnQ-rps16, c) - rps16-trnK, d) - rpl32-trnL.

The nrDNA ITS sequence analysis revealed resolution in the topology of the phylogenetic tree for most species of the genus *Heracleum* (Fig. 5). The pairwise genetic distance between different species ranged from 0 to 2.1%. Meanwhile, for the group of samples including, according to our data, *Heracleum mantegazzianum, H*. cf. *sosnowskyi, H*. cf. *mantegazzianum* and *H*. cf. *sosnowskyi*×*H*. cf. *mantegazzianum* a very low level of variability in the ITS sequence may be noted. The pairwise genetic distance ranged from 0 to 0.3%. A value of 0.3% was associated with a hybrid peak (R) in the 5.8s region in three samples of *H*. cf. *sosnowskyi* collected in the North Caucasus (NCBI: OQ222183, OQ222184, OQ222185). The use of ITS sequences for some representatives of the *Heracleum* genus is known not to lead to their resolution on the topology of the molecular phylogenetic tree in every instance (Logacheva et al., 2010). Based on the variability of ITS sequences, other authors suggested the origin and possible migration routes of 29 species of the *Heracleum* genus in the Chinese flora (Yu et al., 2011).

**Fig. 5.**
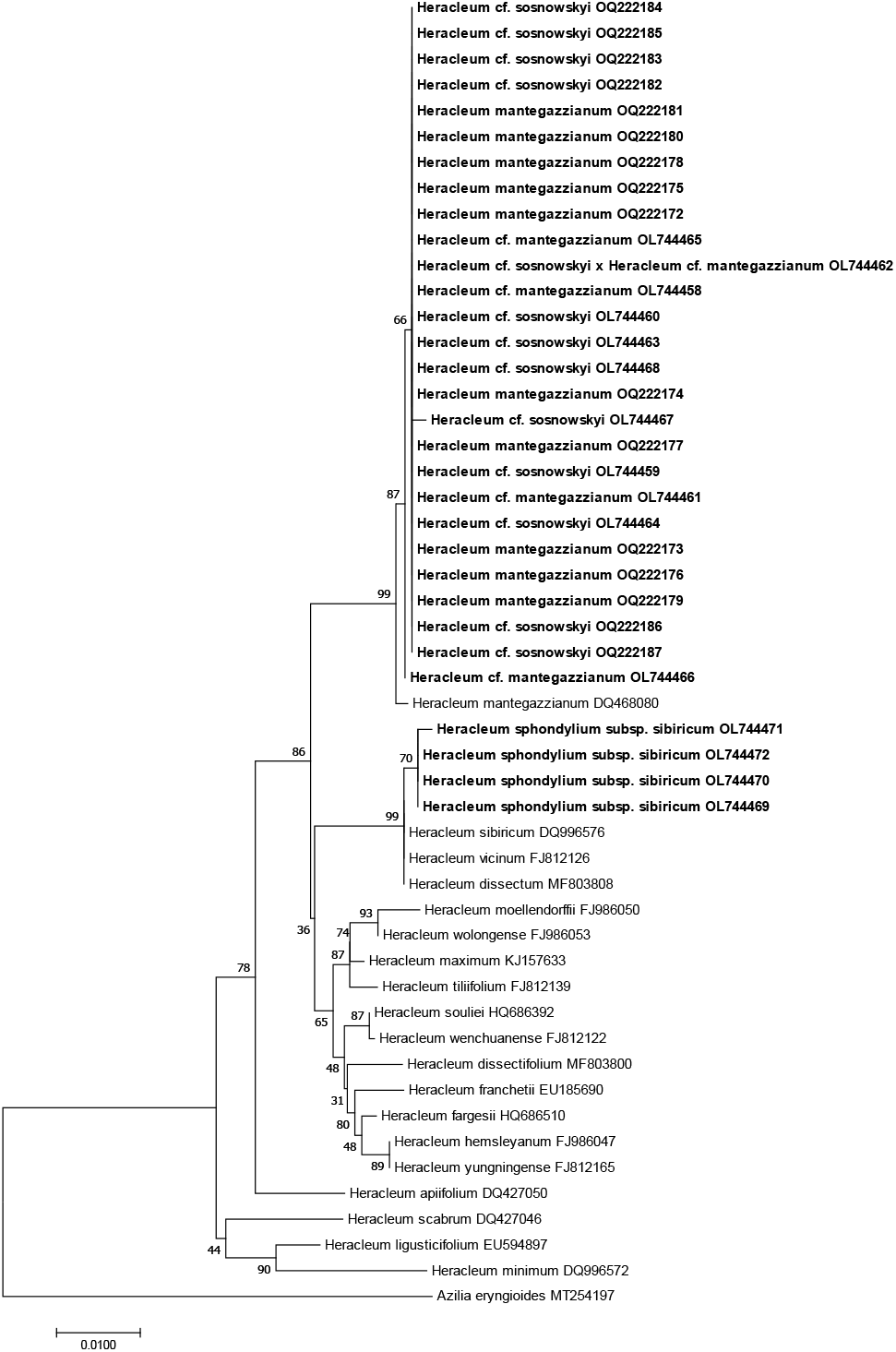
Optimal phylogenetic tree from Maximum Likelihood analyses for the ITS nrDNA *Heracleum* data. The specimen GenBank accession numbers in parentheses follows the names. Samples obtained by the authors are highlighted in bold.

The samples assigned by us to *Heracleum mantegazzianum, H*. cf. *sosnowskyi, H*. cf. *mantegazzianum* and *H*. cf. *sosnowskyi*×*H*. cf. *mantegazzianum* with a high bootstrap value (99) grouped into a separate clade differing from the rest of the species of the genus under (Fig. 5). After overall alignment of the sequences of different species of the *Heracleum* genus at position 125 for ITS1, a unique substitution (A/G) typical for just the mentioned plant samples was found. The sequences we obtained for *H. sphondylium subsp. sibiricum* were found to be identical to the Genbank data for this species (CCЫЛКA), as shown in the molecular phylogenetic tree (Fig. 5). It can be assumed that the ITS sequence has sufficient polymorphism to enable identification of some species of the *Heracleum* genus, but cannot be used to distinguish between the samples of *Heracleum mantegazzianum, H*. cf. *sosnowskyi, H*. cf. *mantegazzianum* and *H*. cf. *sosnowskyi*×*H*. cf. *mantegazzianum* in the European part of Russia.

Thus, using the unique substitution we identified in the nrDNA ITS sequence, we identified a group of samples assigned to *Heracleum mantegazzianum, H*. cf. *sosnowskyi, H*. cf. *mantegazzianum* and *H*. cf. *sosnowskyi*×*H*. cf. *mantegazzianum* from among other representatives of the *Heracleum* genus, with their ITS sequences presented in genetic databases.

Another genetic marker that was used as a potential DNA barcode to distinguish between the analyzed samples was the nrDNA ETS sequence. A revision of genetic databases (NCBI; BOLD Systems) showed that data on ETS are not available for *H. sosnowskyi*, and for *H. mantegazzianum* are presented with regard to one sample.

For 46 samples of plants classified as *Heracleum mantegazzianum, H*. cf. *sosnowskyi, H*. cf. *mantegazzianum* and *H*. cf. *sosnowskyi*×*H*.cf *mantegazzianum* ETS sequences were obtained. A total of 81 ETS sequences for 32 species of the genus *Heracleum* were included in the analysis (Fig. 6). The ratio of variable sites to the total alignment length within the analyzed group (*Heracleum mantegazzianum, H*. cf. *sosnowskyi, H*. cf. *mantegazzianum* and *H*. cf. *sosnowskyi*×*H*. cf. *mantegazzianum*) was 4.2 % (Table 4).

**Fig. 6.**
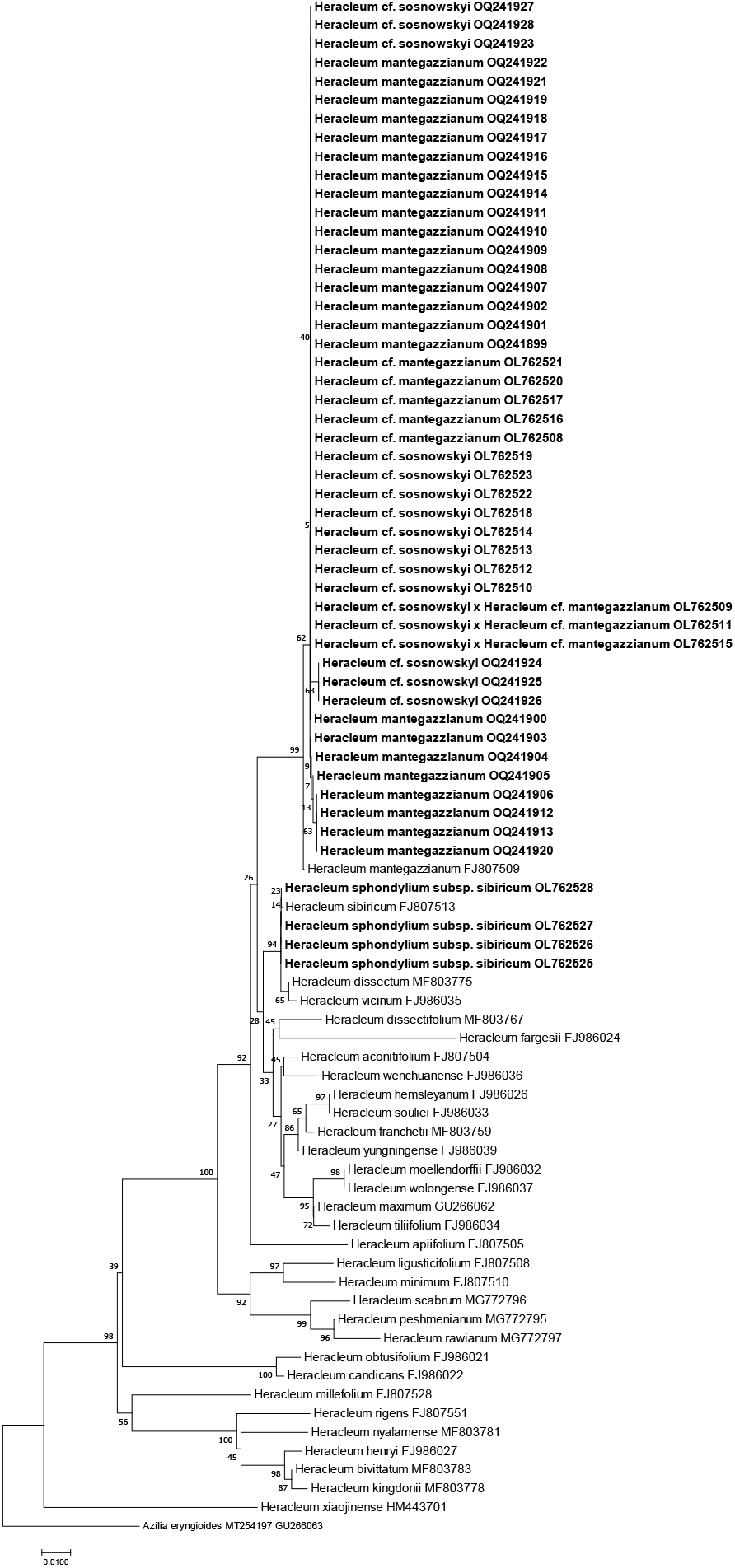
Optimal phylogenetic tree from Maximum Likelihood analyses for the ETS nrDNA *Heracleum* data. The specimen GenBank accession numbers in parentheses follows the names. Samples obtained by the authors are highlighted in bold.

Based on the comparison of *H. sosnowskyi* plant sequences with the only available in genetic databases *H. mantegazzianum* sample, we found the use of the ETS sequence for species distinction upon involving more *H. mantegazzianum* samples as promising (Shadrin et al., 2021). However, the data obtained as a result of follow-up work, analysis of the literature and data on the morphological features of this species group suggest that not all plants previously traditionally assigned to *H. sosnowskyi* based on their confinement to Russia’s European northeast can be attributed specifically to *H. sosnowskyi*.

As a result, the samples that we previously identified as *H. sosnowskyi* (Shadrin et al., 2021) were divided into four groups after reidentification using a set of features (Table 1): *Heracleum mantegazzianum, H*. cf. *sosnowskyi, H*. cf. *mantegazzianum* and *H*. cf. *sosnowskyi*×*H*. cf. *mantegazzianum*.

In addition to the data already obtained (Shadrin et al., 2021), ETS sequences for 24 samples of *H. mantegazzianum* collected from natural populations of the North Caucasus were studied. The qualitative analysis of the ETS sequences of *H. mantegazzianum* from the North Caucasus revealed that they are more similar to plants growing in the European northeast of Russia than with the *H. mantegazzianum* sample presented in the genetic database (NCBI: FJ807509) (Fig. 6). The average pairwise genetic distance within the sample group under analysis ranged from 0.0 to 0.5%. The average pairwise genetic distance between other compared species ranged from 3.0 to 22.7%. A qualitative analysis of sequence variability showed no correlation between the identified changes and the species affiliation of the *H*. cf. *sosnowskyi* and *H. mantegazzianum* samples.

As a result of the analysis, we identified four substitutions at positions 34 (T/C), 186 (A/C, A/G), 259 (C/A) and 357 (A/G) attributable only to 46 samples of the plants classified by us as *Heracleum mantegazzianum, H*. cf. *sosnowskyi, H*. cf. *mantegazzianum* and *H*. cf. *sosnowskyi*×*H*. cf. *mantegazzianum*, including for a sample from the GenBank database (NCBI: FJ807509). The revealed variability can potentially be used as a marker for distinguishing *H. sosnowskyi* and *H. mantegazzianum* from other representatives of the *Heracleum* genus.

The analysis of the alignment of the combined sequences (ITS+ETS = 984 bp) showed the maintained general topology of the phylogenies constructed based on the comparison of the ITS and ETS sequences each separately.

The names *H. mantegazzianum* and *H. sosnowskyi* are likely to be used in one or another part of the vast invasive range of giant hogweed mainly following the established tradition. Obviously, the farms involved in the introduction of a new fodder crop did not carry out any special identification of these plants. Differentiation of two monocarpic species of giant hogweed is not an easy task even for professional botanists, especially since species of the *Heraleum* genus, even belonging to different sections, are capable of crossing (Stewart, Grace, 1984; Jahodová et al., 2007b). The conventionality of the name *H. sosnowsky* can explain the phenomenon of the existence of a clear boundary between the invasive ranges of H. *mantegazzianum* and H. *sosnowskyi:* in the former USSR, the giant invasive species of hogweed is called *H. sosnowskyi*, in Western Europe – H. *mantegazzianum*, in Eastern Europe both species are found. At first glance, from a practical point of view, the subtle taxonomic differences between the two species seem not to matter. However, misidentification of invasive species is scientifically unacceptable and can have serious implications in practice. The name *H. sosnowskyi* was included in official and unofficial methodological guidelines for the eradication of unwanted thickets of this species, in legal acts on regional and municipal levels in Russia, Belarus, Latvia, Lithuania and Estonia.

## Conclusion

The comparative analysis of the following sequences: *rbc*L, *mat*K, rps16 intron, cpDNA intergenic spacers *psb*A-*trn*H, *trn*Q-*rps*16, *rps*16-*trn*K and *rpl*32-*trn*L for the samples defined by us as *Heracleum mantegazzianum, H*. cf. *sosnowskyi, H*. cf. *mantegazzianum* and *H*. cf. *sosnowskyi* × *H*. cf. *mantegazzianum* did not reveal any qualitative differences that could be correlated with the samples. The comparative analysis of the nrDNA ITS and ETS sequences revealed qualitative differences in them, which make it possible to identify the groups of samples: *Heracleum mantegazzianum, H*. cf. *sosnowskyi, H*. cf. *mantegazzianum* and *H*. cf. *sosnowskyi* × *H*. cf. *mantegazzianum* among other representatives of the *Heracleum* genus, which are referred to in genetic databases at the moment. The comparison of ITS and ETS sequences for the samples of *Heracleum mantegazzianum, H*. cf. *sosnowskyi, H*. cf. *mantegazzianum* and *H*. cf. *sosnowskyi* × *H*. cf. *mantegazzianum* showed no correlation between their polymorphism and the type of samples in the given sample series. The data obtained through the study, as well as information on morphological features that make it possible to distinguish between *H. mantegazzianum* and *H. sosnowskyi* indicate that it is possible that the names *H. mantegazzianum* and *H. sosnowskyi* are used in one or another part of the vast invasive range of giant hogweed mainly following the established tradition.

